# Abatacept treatment in LRBA deficient patients results in an increase in circulating FoxP3^+^Helios^+^ natural Treg

**DOI:** 10.1101/2024.02.10.579765

**Authors:** Sabine Donhauser, Emilia Salzmann-Manrique, Leon Maximilian Lueck, Julia Fekadu-Siebald, Christoph Königs, Ralf Schubert, Celia Kaffenberger, Laura M. Moser, Peter Bader, Sabine Huenecke, Shahrzad Bakhtiar

## Abstract

**Background:** Mutations in the lipopolysaccharide-responsive beige-like anchor (LRBA) gene were identified to cause autoimmunity and immunodeficiency with a broad spectrum of clinical manifestations. Increased lysosomal degradation of the CTLA-4 due to the lack of recylcing by LRBA protein has been described resulting in a reduced suppressive capacity of regulatory T cells (Treg).

**>Objective:** We sought to explore the extended Treg profile of patients with LRBA deficiency (N=6) compared to reference values for Treg subpopulations from an age-matched healthy cohort (N=39) prior and during abatacept treatment as well as after allogeneic haematopoietic stem cell transplantation. In parallel we assessed the feasibility and robustness of the CHAI and the IDDA2.1 scores for this cohort.

**Methods:** Using a flow cytometric approach with a pre-formulated antibody panel in peripheral blood samples, we analyzed Treg subsets including CD4^+^CD25^hi^FoxP3^+^ Treg, Helios^+^ natural Treg, Helios^-^ induced Treg, CD39^+^ Treg, CD62L^+^CD45RA^+^ naïve Treg, CD62L^+^CD45RA^-^ memory Treg, and FoxP3^hi^CD45RA^-^ effector Treg as well as CD4^+^CD25^hi^CD127^low^ Treg. Longitudinal data were collected while patients were receiving abatacept and in three patients after alloHSCT. CHAI and IDDA2.1 scores were performed.

**Results:** The healthy cohort including individuals from the ages of 2 to 40 years showed a steady total CD25^hi^FoxP3^+^ Treg population around 5%, while there was a significant age-dependent increase in Helios^+^ natural Treg (P=0.003), Helios^-^ induced Treg (P=0.046), CD62L^+^CD45RA^-^ memory Treg (P=0.020) with a continuous decrease in CD62L^+^CD45RA^+^ naïve Treg (P=0.015) and CD4^+^CD25^hi^CD127^low^ Treg (P=0.024) over time. LRBA deficient patients showed a significant lack in FoxP3^+^Helios^+^ natural Treg (P=0.003), CD62L^+^CD45RA^+^ naïve Treg (P<0.001), FoxP3^hi^ CD45RA^+^ effector Treg (P=0.001) and CD4^+^CD25^hi^CD127^low^ Treg (P=0.005), while their CD62L^+^CD45RA^+^ memory Treg (P=0.016) were significantly elevated. Abatacept treatment led to a significant increase in natural Treg (P=0.003) in patients without having a measurable effect on the other subpopulations. This was accompanied by a decrease in sIL2R levels. The IDDA2.1 score was feasible for this mainly pediatric cohort and correlated with patients’ clinical course of the disease.

**Conclusion:** Within the Treg subsets in peripheral blood of LRBA deficient patients, there is a significant lack of natural, naïve, effector and CD4^+^CD25^hi^CD127^low^ Treg, while memory Treg are elevated. During abatacept treatment we observe a significant increase in circulating Helios^+^ natural Treg levels. IDDA2.1 scoring system is feasible for pediatric patients to assess the severity of their disease and the response to the treatment.

## 1. Introduction

Regulatory T cells (Treg) are understood as main key players in maintaining the peripheral immune tolerance. After their initial description [1], there has been growing knowledge around Treg subsets and their subset specific mechanisms of action. A milestone in understanding Treg was the discovery of the function of Forkhead Box P3 (FoxP3) [2]. [3].

The currently accepted concepts describe Treg based on their development: thymic derived ‘natural’ Treg (nTreg) and induced Treg (iTreg). It was shown that nTregs express both effector molecules, including CTLA-4, LAG-3, CD39 and CD73, as well as CD28, CD80/86 (B7), CD40, OX40 and 4-1BB with co-stimulatory function. [4], [5], [6] iTreg are induced from CD4+ T cells by antigenic stimulation and other factors such as inflammation, tumor and infection. [7].

The knowledge of expression of IL-7 receptor alpha (CD127) allowed to define the population of CD4^+^CD25^hi^CD127^low^ supposed to be correlating to the nTreg [8] which became a routine strategy to assess Treg in immune diseases. Helios, an Ikaros family transcription factor, has also been associated to the nTreg with somewhat contradicting data. [9], [10].

Despite differences in nomenclature and description of their surface markers, the following Treg subsets have been assigned to the circulating Treg in peripheral blood of humans: CD4^+^FoxP3^+^ Treg, Helios^+^ natural Treg, Helios^-^ induced Treg, CD39^+^ Treg, CD62L^+^CD45RA^+^ naïve Treg, CD62L^+^CD45RA^-^ memory Treg, and FoxP3^hi^CD45RA^-^ effector Treg. [10], [11], [12], [13] (schematic overview given in Figure 1 A+B).

**Figure 1:**
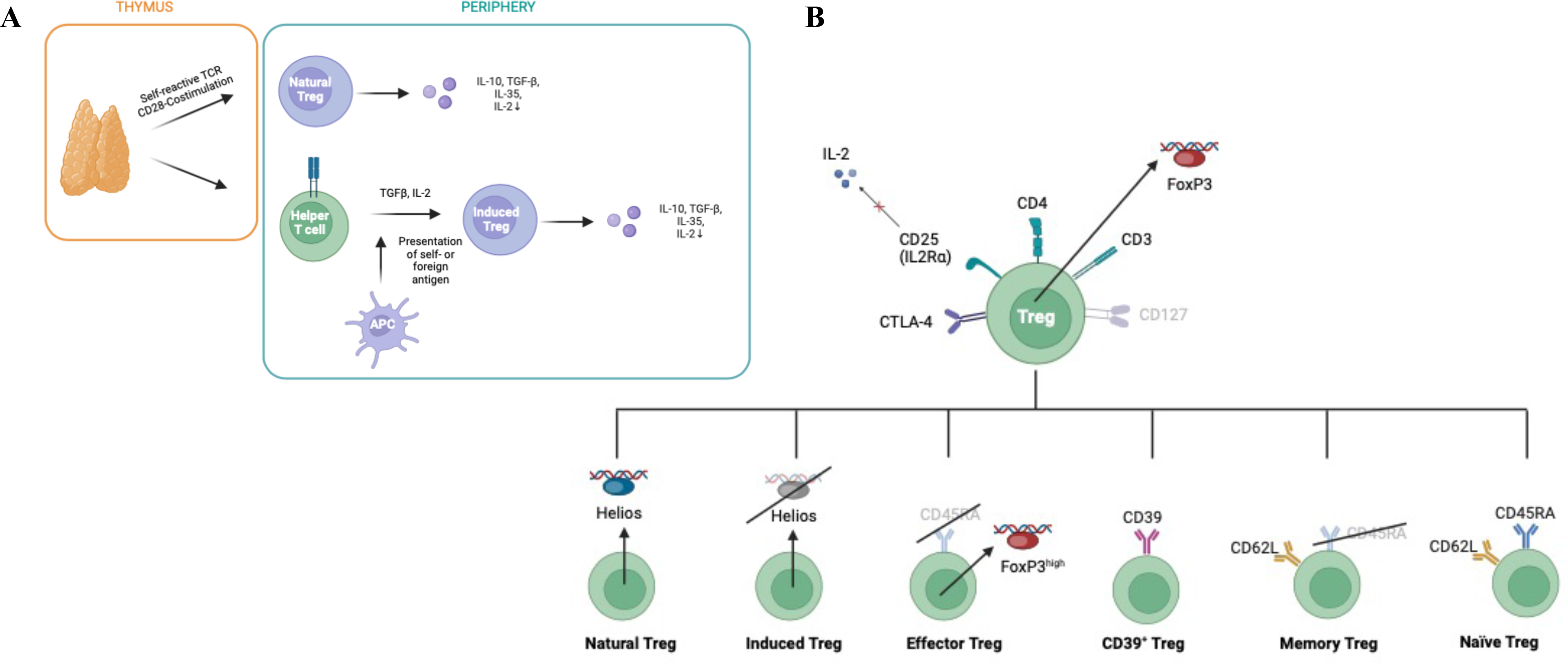
Schematic overview using Biorender.com. Natural Treg originate in the thymus while induced Treg can arise from conventional T cells triggered by antigen presenting cells or stimulated by cytokines such as TGF-β or IL-2. Both species produce anti-inflammatory acting cytokines including TGF-β, IL-10, IL-35 and also suppress the amount of circulating proinflammatory IL-2. APC, antigen presenting cell; TGFβ, Transforming growth factor β; IL-, Interleukin. Figure modified from [10], [11] and [12]. Illustration of the Duraclone® antibody panel for phenotyping Treg subsets via their surface and intracellular markers with additional use of anti-CD62L. Treg are defined through their transcription factor FoxP3, the constitutive expression of CTLA-4 and CD25 on the cell surface as well as the low expression of CD127. In this model Helios is a marker for natural Treg, while induced Treg are defined as Helios^-^. CD39 expression among Helios^+^ Treg characterizes a further Treg subgroup. Effector Treg are distinguished by high expression level for FoxP3. Naïve Treg differ from memory Treg in the expression of CD45RA. FoxP3, Forkhead-Box-Protein P3; CTLA-4, cytotoxic T-lymphocyte-associated Protein 4. Figure modified from [10], [12] and Duraclone® Treg Panel (Beckman Coulter).

One major step in understanding the role of Treg in human disease was the evidence that mutations of the human FoxP3 gene cause a severe disease of immune dysregulation, polyendocrinopathy, enteropathy, X-linked syndrome, called IPEX syndrome. [14],[15] During the tremendous progress in the field of monogenetic inborn error of immunity (IEI) a series of IPEX-like diseases have been described, summarized in the current IUIS classification. [16] Mutations in the lipopolysaccharide-responsive beige-like anchor (LRBA) gene were identified to cause progressive immune dysregulation with poor quality of life and high mortality without genotype phenotype correlation, which is a prime example for this disease category. [17] [18] LRBA prevents CTLA-4 degradation in endocytic lysosome resulting in a disturbed suppressive capacity of Treg in case of LRBA deficiency. [19] Although LRBA has been shown in colocalization with a series of other proteins, there has been no other cargo found. [20] Trying to restore this disbalance, treatment with CTLA-4 fusion protein (abatacept), which binds to CD80/CD86 (B7) on APC, is the an effective modulatory treatment so far available for the affected patients. [21] For severely affected patients allogeneic stem cell transplantation (alloHSCT) is the ultimate curative option. [22], [23].

Regarding immune cell subsets, there has been evidence for decreased expression of FoxP3, CD25, Helios and CTLA-4 on Treg in LRBA deficiency patients. [24] Their T-cell repertoire seems to be skewed in favor of memory T cells with marked expansion of T follicular helper and contraction of T follicular regulatory cells with normal frequencies of induced Treg cells. They also show high levels of autoantibody production. [19] We recently reported a disturbed maturation of regulatory B cells with an increased expansion of autoimmunity related CD21^low^ expressing B cells, which significantly decreased while patients were receiving abatacept. [25].

In this work we provide an extended analysis of Treg subsets with a focus on Helios^+^ natural Treg and Treg subsets while patients were receiving abatacept to assess whether disease treatment influences circulating Treg. Reference values of peripheral blood Treg subsets according to age was developed allowing an age-matched analysis between healthy and the LRBA cohort. Disease symptoms were scored by IDDA2.1 score at treatment start and during the longitudinal analysis while patients were receiving abatacept. [26].

A detailed analysis of Treg subsets can contribute to the understanding of the heterogeneity of clinical symptoms in this disease. These Treg markers might provide disease specific targets for future immunotherapies.

## 2. Patients and Methods

### 2.1 Study cohort

This study was conducted between June 2020 and August 2023. We recruited six patients with LRBA deficiency syndrome at our center in Frankfurt in Germany who had not previously been treated with abatacept. The study protocol was approved by the ethics committee of the Goethe University Frankfurt am Main (IRB approval, Ref. No. 436/16). All the patients and parents signed an informed consent form in accordance with the Declaration of Helsinki.

Patients’ ages ranged from 3.4 to 24.2 years (median: 14.5 years). All patients suffered from symptoms of immune dysregulation and autoimmunity with symptom variability. The major clinical problem in this cohort was enteropathy and associated malnutrition and failure to thrive, followed by autoimmune cytopenia and need for immunoglobulin substitution in all patients. In addition, all patients had splenomegaly and most of them showed lymphadenopathy and skin manifestations such as eczema, alopecia and vitiligo. Four patients were affected by endocrinopathy which manifested as autoimmune thyroiditis or insulin-dependent diabetes mellitus type 1. Half of the cohort had chronic or recurrent infections. Granulomatous lymphocytic interstitial lung disease and occurrence of a malignancy were each observed in one patient. Arthritis, pancreatitis and nephropathy also occurred in some patients, but were seen less frequently. (Table 1).

**Table 1:**
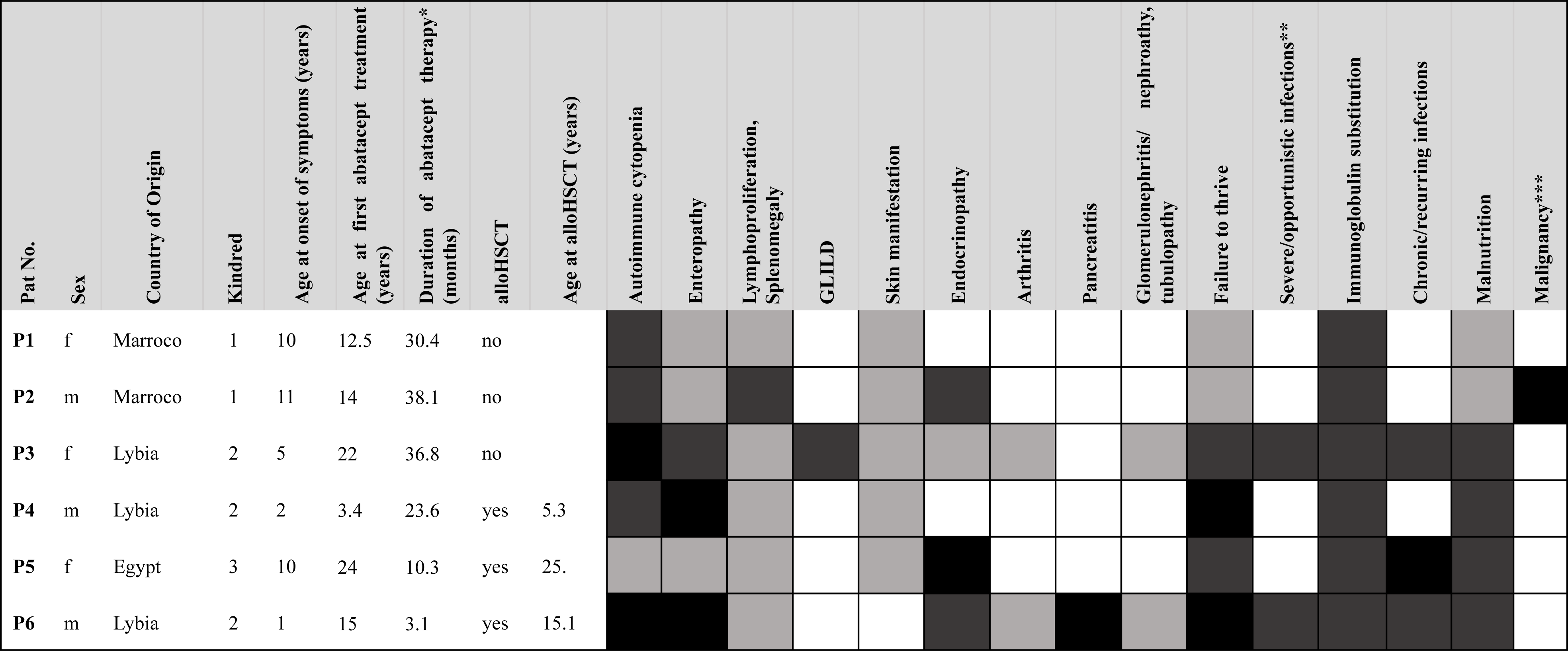
Patient characteristics showing patients’ gender and country of origin, age at first symptoms, related disease symptoms before start treatment, age at abatacept start and alloHSCT procedure. The disease symptoms were classified according to the IDDA2.1 score. Shading based on IDDA2.1 Score: white = 0 points (absent), light grey = 1 or 2 points (mild or moderate, not requiring treatment or intermittent therapy needed), dark grey = 3 points (severe, continuous therapy needed), black = 4 points (life-threatening, refractory, irreversible) *excl. chronic infections, **black shading: yes, white shading: no, ***P1-3 currently still on abatacept, duration in months until October 27, 2023. GLILD: Granulomatous–lymphocytic interstitial lung disease.

Additionally, peripheral blood for routine laboratory examination and clinical data of the patients were collected for immune phenotyping at first examination as well as after biweekly abatacept treatment (10–15 mg/kg i.v.) and following successful alloHSCT (N=3).

Reference values for Treg subsets were obtained in a healthy cohort including 39 children and adults who did not have any chronic diseases or infections and were not under medication. The adult controls were healthy volunteers. For individuals below the age of 18 years, residual blood samples following blood tests for the routine assessments were used.

### 2.2 Sample preparation and flow cytometric analysis

Whole blood samples were collected and analyzed within 24 hours of collection. Following incubation with separately added anti-CD62L, the cells were washed with PBS once, treated with FCS and then fixed and permeabilized using PerFix reagent buffers (PerFix-nc Kit, Beckman Coulter). Afterwards they were incubated in pre-formulated tubes (DuraClone®; Beckman Coulter) for surface and intracellular staining followed by a further washing step with PBS. Staining for CD127 and CD25 to rule out CD25^hi^CD127^low^ Treg was performed separately in the clinical laboratory routine with anti-CD25 and anti-CD127 incubation performed first, followed by erythrocyte lysis. Analyses were performed using a ten-color flow cytometer (Navios, Beckman Coulter, Krefeld, Germany) and five-color flow cytometer (Cytomics FC500, Beckman Coulter, Krefeld, Germany). The Gating was carried out with the Kaluza Analysis software (Kaluza Analysis 3.1, Beckman Coulter, Krefeld, Germany). First, in the CD45 vs SS dot plot a gate was placed on the lymphocytes (Fig. 2A) which were then gated on CD3^+^CD4^+^ T cells (Fig. 2B). Treg were gated as CD25^hi^FoxP3^+^ (Fig. 2C) and as CD25^hi^ (Fig. 2D), the latter being subgrouped in E-H. The gate FoxP3 vs. Helios allowed the distinction between natural Treg (FoxP3^+^Helios^+^) and induced Treg (FoxP3^+^ Helios^-^) (Fig. 2E) and the Helios vs. CD39 gate shows the CD39^+^ cells (Fig. 2F). The expression level of CD45RA makes the classification into naïve (CD62L^+^CD45RA^+^) and memory Treg (CD62L^+^CD45RA^-^) possible (Fig. 2G). To rule out non Treg, effector Treg were determined to be highly positive for FoxP3 and not expressing CD45RA (Fig. 2H). (Figure 2 A+H).

**Figure 2:**
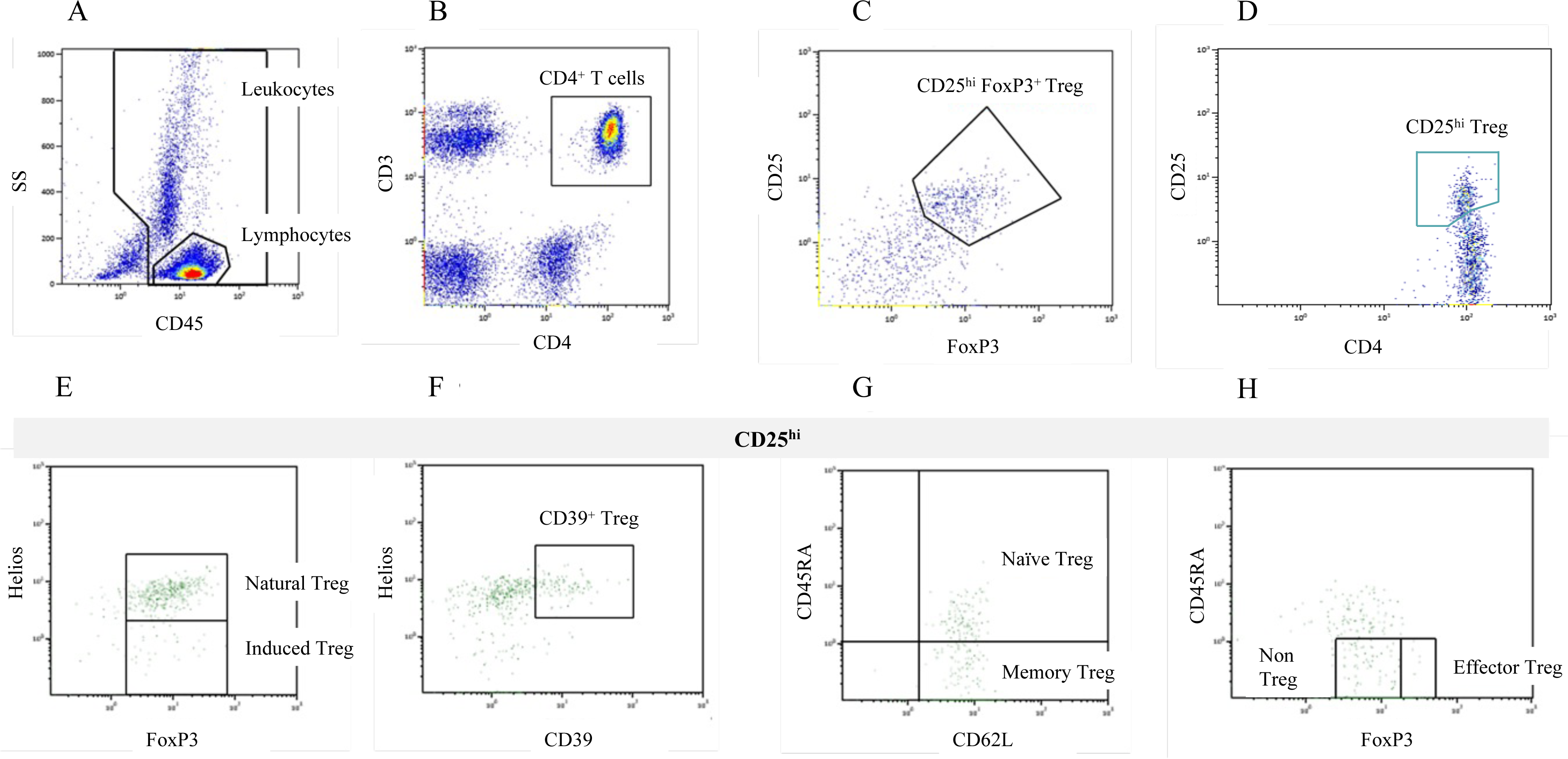
Gating strategy for Treg (%) subpopulation analysis. Flow cytometric analysis was performed on whole blood samples after antibody labeling with the Duraclone® Treg Panel and additional Anti-CD62L antibody. First in the CD45 vs SS dot plot a gate was placed on the lymphocytes (A) which were then gated on CD3^+^CD4^+^ T cells (B). The main Treg population was gated as CD25^hi^FoxP3^+^ (C) and as CD25^hi^ (D), the latter being subgrouped in E-H. The gate FoxP3 vs. Helios allows the distinction between natural Treg (Helios^+^FoxP3^+^) and induced Treg (Helios^-^ FoxP3^+^) (E) and the Helios vs. CD39 gate shows the Helios^+^CD39^+^ cells (F). The expression level of CD45RA allows the classification into naïve (CD62L^+^CD45RA^+^) and memory Treg (CD62L^+^CD45RA^-^) (G). To rule out non Treg, effector Treg were determined to be highly positive for FoxP3 and not expressing CD45RA (H). SS, side scatter.

### 2.3 Clinical score for diseases with immune dysregulation

For prospective monitoring of diseases with immune dysregulation, we used the two currently available scores; The Immune Deficiency and Dysregulation Activity (IDDA 2.1) [26] and the CHAI-Morbidity [27] scores. Because of the mainly pediatric cohort, the CHAI-Morbidity score could not be performed with all details. Especially repetitive lung function tests and chest CT scans were not feasible in young children. Data set could be completed for the IDDA2.1 score comprising 22 parameters on a 2–5-step scale. 2–5-step scale.

### 2.4 Statistical analysis

A reference model of the relative frequencies (% of CD4^+^ T helper or % of CD25^hi^ T cells) for each Treg subpopulation was performed with the values from samples of healthy donors. This regression model used cubic B-splines, allowing for a possible nonlinear relationship between age and the relative distribution of cells in each subpopulation. Differences in the percentage distribution between patients and healthy individuals were analyzed by t-test for paired observations. We compared the relative frequency values from the patients’ first measurement, achieved before the start of treatment, with the corresponding age-matched expected mean from the reference model. To evaluate the effect of abatacept on Treg subsets over time, B-spline mixed effect regression model was used. To take age dependency into account, each relative frequency value was normalized to the corresponding age-specific expected mean reference value. Then, each Treg subsets were analysed considering time as fixed effect and patient as random effect. All analyses were performed with the use of R statistical computing environment software version 4.03 (R Foundation for Statistical Computing, Vienna, Austria). All statistical testing was performed with the use of two-tailed tests; a P value of less than 0.05 was considered to indicate statistical significance.

## 3. Results

### 3.1 Age-matched reference model shows an age-dependent maturation of Treg subsets in healthy controls

The healthy cohort including individuals from 2 to 40 years of age showed a steady total FoxP3^+^ Treg population around 5% (3B), while there was a significant age-dependent increase in Helios^+^ natural Treg (P=0.003) (3C), Helios^-^ induced Treg (P= 0.046) (4D), CD62L^+^CD45RA^-^ memory Treg (P=0.020) (3G) with a continuous decrease in CD62L^+^CD45RA^+^ naïve Treg (P=0.015) (3F) and CD4^+^CD25^hi^CD127^low^ (P= 0.024) (3 I) by the age of 40 years. The predicted reference values (%) of Treg subsets, with their 95% confidence interval, are presented for representative ages in Suppl. Table 1.

### 3.2 LRBA deficient patients show a significant lack in circulating naïve Treg and natural Treg while having normal induced Treg

The first Treg subsets measurements, previously any treatment, was compared to its corresponding age-matched mean reference value. The figure 4 exihibts the values for the patient cohort with respect the reference curves. Detailled summary with median (range), mean, standard desviation (SD), paired mean difference and its 95% confidence intervals are presented in Table 2.

**Figure 3:**
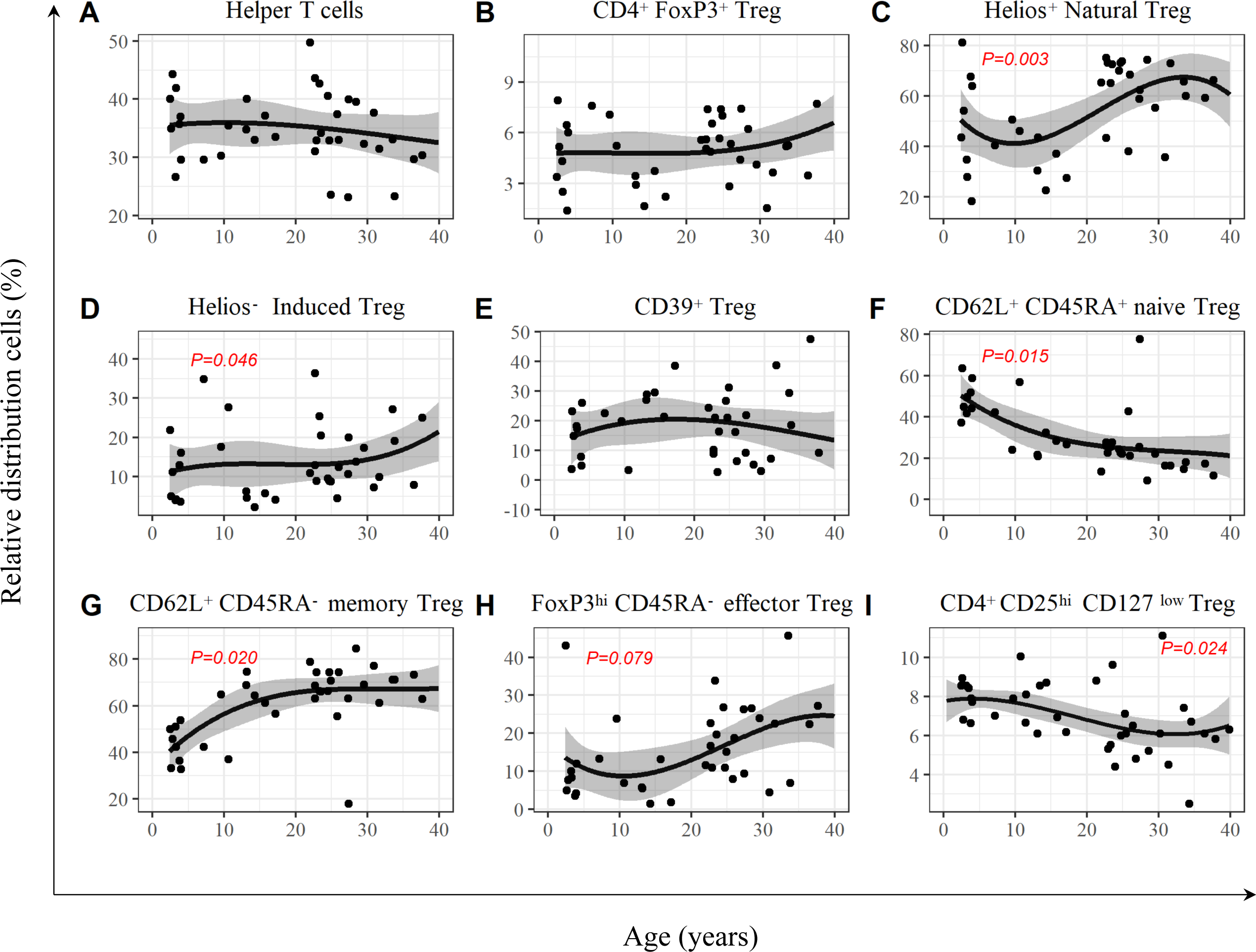
Age-dependent development of T helper cells, Treg and their subsets in 39 healthy volunteers. The black curve showes the estimated mean; the grey shaded area illustrated the 95% confidence interval for the T helper cells (CD3^+^CD4^+^ T cells) (A) and FoxP3^+^Treg (CD25^hi^FoxP3^+^) (B), as well as the subsets: Natural Treg (Helios^+^FoxP3^+^) (C), induced Treg (Helios^-^FoxP3^+^) (D), CD39^+^ Treg (Helios^+^CD39^+^) (E), naïve Treg (CD62L^+^CD45RA^+^) (F), memory Treg (CD62L^+^CD45RA^-^) (G), effector Treg (FoxP3^hi^CD45RA^-^) (H) and CD25^hi^CD127^low^ Treg (I). On the y-axis, the values are shown as the percentage of lymphocytes (for A) and T helper cells (for B) or CD25^hi^ Treg (for C-I). The x-axis shows the age in years. P-values of the statistical analysis of the age dependency of the distribution that show a tendency (P<0.1) or show significance (P<0.05) are shown in red letters. P values from linear regression model determined using student’s t-test for coefficients.

**Figure 4:**
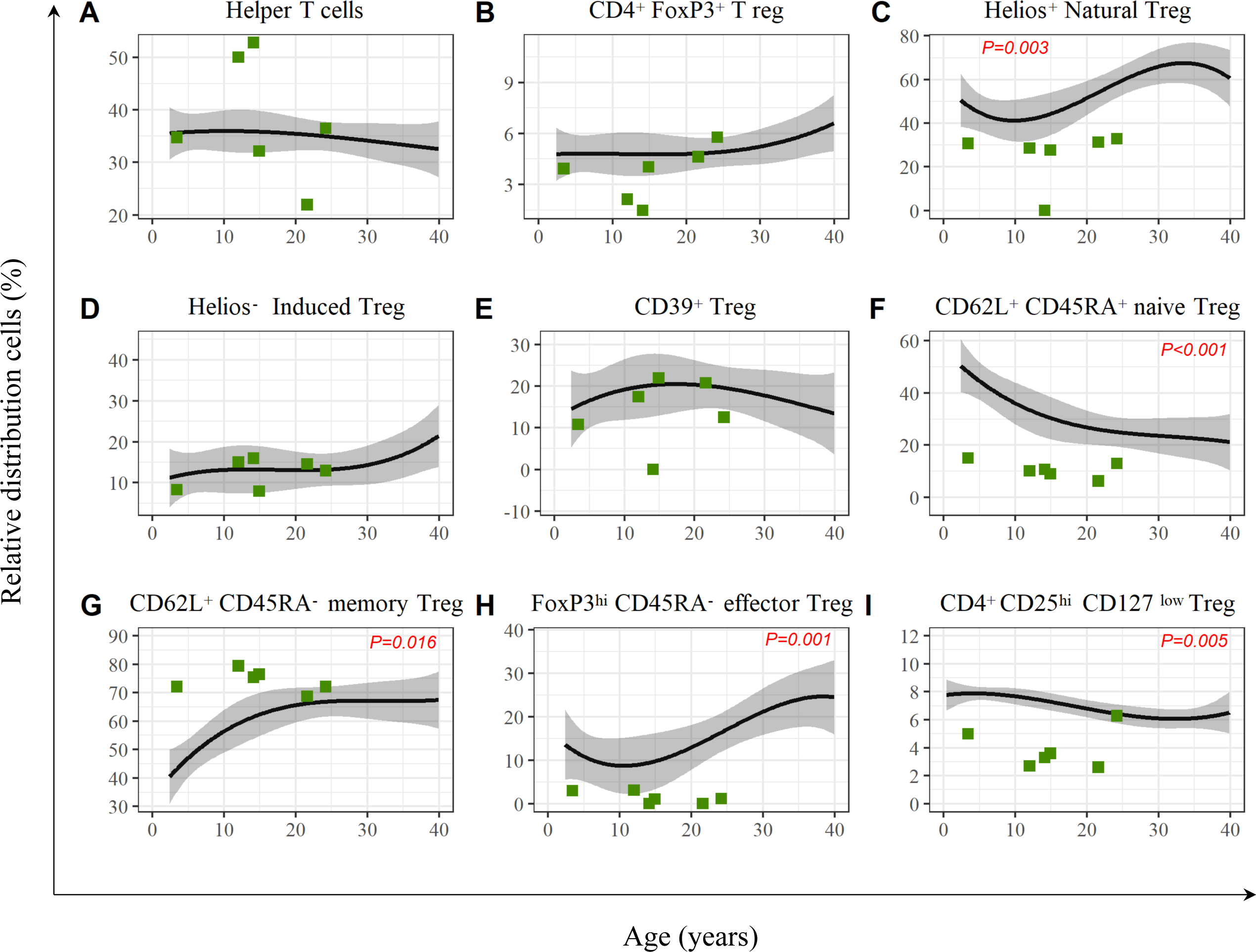
Representation of the relative distribution of T helper cells, Treg and their subpopulations in the patient cohort with LRBA deficiency compared to the in Figure 3 described distribution in the healthy cohort. We observe a significant lack of natural Treg (p=0.003), naïve Treg (p<0.001), effector Treg (p=0.001) and CD25^hi^CD127^low^ Treg (p=0.005) and a slight excess in memory Treg (P=0.016) in the LRBA deficient cohort. P value determined using two-tailed paired t-test.

**Table 2:** Relative values of conventional T helper cells and regulatory T cells and their subtypes. The median, range, mean (%), the standard deviation (%) as well as the paired difference with 95% confidence interval and the P values are given for each Treg subset comparing LRBA deficient patients to their age-matched cohort of healthy subjects. While there is no difference in CD4^+^ T cell population and FoxP3^+^ Treg, there is a significant lack seen in patients for natural Treg (P=0.003), naïve Treg (P<0.001), and effector Treg (P=0.001), while patients seem to have slightly elevated memory Treg (P=0.016). CD4^+^CD25^hi^CD127^low^ Treg population shows the same pattern as natural or naïve Treg. FoxP3, Forkhead-Box-Protein P3, SD standard deviation, CI confidence interval. P value determined using two-tailed paired t-test.

|  | Median (range) % |  | Mean (SD) % |  | Paired Mean difference% (95% CI) | P value |
| --- | --- | --- | --- | --- | --- | --- |
|  | Age-matched healthy cohort | LRBA cohort | Age-matched healthy cohort | LRBA cohort |  |  |
| CD4<sup>+</sup> T cells | 35.7 ( 35.0 to 35.9) | 35.6 (21.9 to 52.8) | 35.6 ( 0.4) | 38.0 (11.6) | 2.4 (-9.5 to 14.3) | 0.621 |
| FoxP3<sup>+</sup> regulatory T cells | 4.8 (4.8 to 4.9) | 4.0 (1.5 to 7.6) | 4.8 (0.1) | 4.0 (1.6) | -1.1 (-2.8 to 0.5) | 0.134 |
| Natural regulatory T cells | 46.1 (41.8 to 58.5) | 29.6 (0 to 32.4) | 48.4 (6.7) | 25.2 (12.5) | -23.2 (-34.6 to -11.8) | 0.003 |
| Induced regulatory T cells | 13.0 (11.6 to 13.2) | 13.8 (7.9 to 16.0) | 12.9 (0.6) | 12.5 (3.5) | -0.5 (-3.8 to 2.9) | 0.744 |
| CD39<sup>+</sup> regulatory T cells | 20.0 (15.4 to 20.4) | 15.0 (0 to 22.0) | 19.3 (1.9) | 14.0 (8.1) | -5.3 (-13.7 to 3.1) | 0.164 |
| Naïve regulatory T cells | 30.8 (25 to 48.1) | 10.4 (6.3 to 15.1) | 32.4 (8.4) | 10.7 (3.1) | -21.7 (-28.8 to -14.6) | <0.001 |
| Memory regulatory T cells | 61.9 (43.1 to 66.8) | 73.8 (68.8 to 79.5) | 59.9 (8.7) | 74.1 (3.8) | 14.2 (4.0 to 24.4) | 0.016 |
| Effector regulatory T cells | 11.1 (8.8 to 16.5) | 1.0 (0 to 3.1) | 11.9 (3.1) | 1.4 (1.4) | -10.5 (-14.3 to -6.7) | 0.001 |
| CD4<sup>+</sup>CD25<sup>hi</sup>CD127<sup>low</sup> regulatory T cells | 7.3 (6.4 to 7.9) | 3.5 (2.6 to 6.3) | 7.2 (0.6) | 3.9 (1.5) | -3.3 (-5.0 to -1.5) | 0.005 |

Compared to their age-matched control cohort the LRBA deficient cohort showed no significant alterations in the leukocytes and CD4^+^ lymphocytes, which were 35.6±0.4% (mean±SD, median 35.7%) in the age-matched healthy cohort versus 38.0±11.6% (median 35.6%) for the LRBA deficient cohort (P=0.621). (Figure 4A) The analysis of Treg cell subsets is given in Figure 4 (B-I) as follows:

**FoxP3^+^ Treg** were stable at 4.8 ±0.1% (median 4.8%)) in the age-matched healthy cohort versus 4±1.6% (median 4.0%) in patient cohort; P=0.134 (Figure 4B).

**Helios^+^ natural Treg** were found in 48.4±6.7% (median 46.1%) in the age-matched healthy cohort, while LRBA deficient patients showed a homogenous pattern with a significantly reduced level in all affected individuals (mean: 25.2±12.5% (median 29.6%)); P=0.003 (Figure 4C).

**Helios^-^ induced Treg** were detected at a level of 12.9±0.6% (median 13.0%) in the age-matched healthy cohort. All LRBA deficient patients were found to have their induced Treg within the reference range (mean: 12.5±3.5% (median 13.8%)); P=0.744 (Figure 4D).

**CD39^+^ Treg** were at 19.3±1.9% (median 20%) in the age-matched healthy cohort versus 14±8.1% (median 15.0%) for the LRBA deficient cohort; P=0.164 (Figure 4E).

**CD62L^+^CD45RA^-^ naïve Treg** were found at 32.4±8.4% (median 30.8%) within the healthy cohort, while the patients cohort showed significantly reduced levels of 10.7±3.1% (median 10.4%). The mean difference in the paired observations was −21.7% (95% CI −28.8 to −14.6%) P < 0.001 (Figure 4F).

**CD62L^+^CD45RA^-^ memory Treg** were at 59.9±8.7% (median 59.9%) within the healthy cohort, while the patients’ cohort were found to have significantly higher levels 74.1±3.8 (median 73.8%); P=0.016 (Figure 4G).

**FoxP3^hi^CD45RA^-^ effector Treg** were at 11.9±3.1% (median 11.1%) within the healthy cohort, while the patients’ cohort were found to have significantly lower, almost undetectable levels, in mean 1.4±1.4% (median 1.0%); P=0.001 (Figure 4H).

**CD25^hi^CD127^low^ Treg** were at 7.2±0.6% (median 7.3%) within the healthy cohort, while the patient cohort were found to have significantly lower levels, in mean 3.9±1.5% (median 3.5%); P=0.005 (Figure 4I).

### 3.3 Biweekly intravenous abatacept increases circulating Helios^+^ natural Treg

Treg subsets were analyzed prior, at the time of abatacept start and following treatment in each patient. Interestingly, we observed that two to four weeks after abatacept treatment start, natural Treg percentages decreased to a level clearly below the starting point in all patients beside P3, whose first post abatacept measurement was 4.5 months post treatment start. After reaching the lowest point, natural Treg significantly increased to the level of age-matched reference range in P1, P2, P3, P4 and P5. Patient 6 received an early alloHSCT and was assessed post-transplant. (Figure 5). Therapy with abatacept seems to result in a increase of natural Treg after the two initials months (P <0.002). Treg subsets before and during abatacept treatment including following alloHSCT in P4, P5, P6 are shown in Suppl. Figure 1. Post-transplantation there was an early immune reconstitution for Treg subsets.

**Figure 5:**
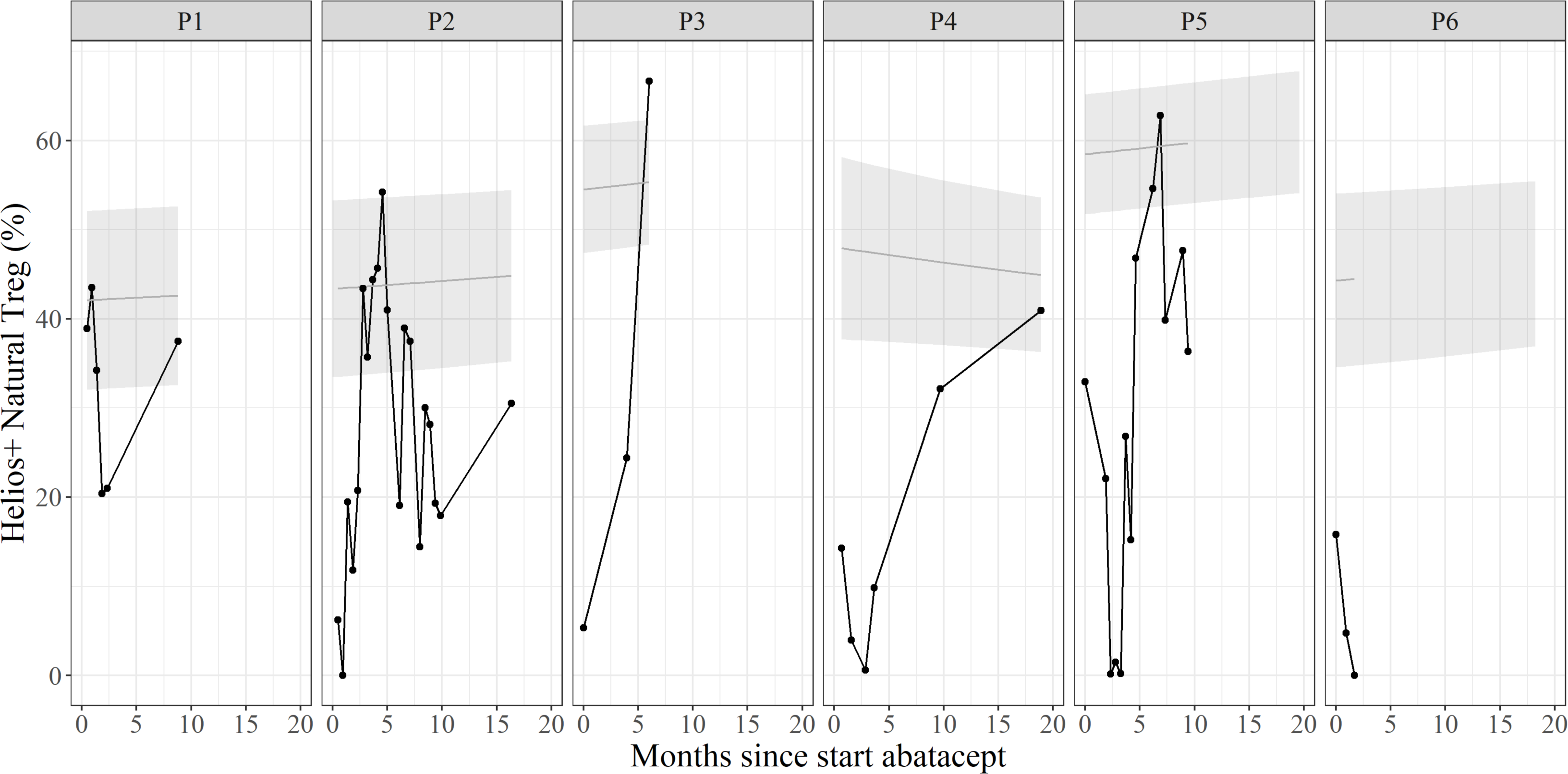
Longitudinal development of the relative distribution of natural Treg (%) in the patient cohort with LRBA deficiency while receiving abatacept compared to the in Figure 3 described healthy controls. Two to four weeks after abatacept treatment start, natural Treg counts decreased to a level below the starting point in all patients beside P5, whose first post abatacept measurement was 4.5 months post treatment start. After reaching the lowest point, natural Treg significantly increased to the level of normal in P1, P2, P3, P4 and P5. Patient 6 received an alloHSCT and was assessed afterwards.

### 3.4 Clinical Score using IDDA2.1 score

The disease severity was assessed by the responsible physician at each visit. All 22 items of IDDA2.1 score could be assessed for the patients. One patient suffered from a malignancy previously (P2), which is an additional item in the IDDA2.1 scoring system. Severe HLH was not observed in this cohort. The highest starting point was 52 for P3, which is clearly the sickest patients in this cohort. The IDDA2.1 score increased in this patient to >70 points despite being partially responsive to abatacept. All other patients decreased their score significantly, in keeping with improvement of their clinical status and laboratory data. Those patients receiving an alloHSCT improved almost entirely. The score dropped to < 10 points.

Taken together, the intestinal symptoms could be well controlled, autoimmune gastritis and colitis were regressive and chronic diarrhea was improved. In two patients however, intestinal symptoms were progressive (P3, P5). Autoimmune cytopenias responded well to the treatment in all patients. Skin manifestations remained largely unchanged, at most one patient showed a lack of recurrence of aphthae and others a slight improvement in skin eczema. IDDM and autoimmune thyroiditis remained unaffected as well as hypogammaglobulinemia so that all patients still needed substitution. (Figure 6A).

**Figure 6A:**
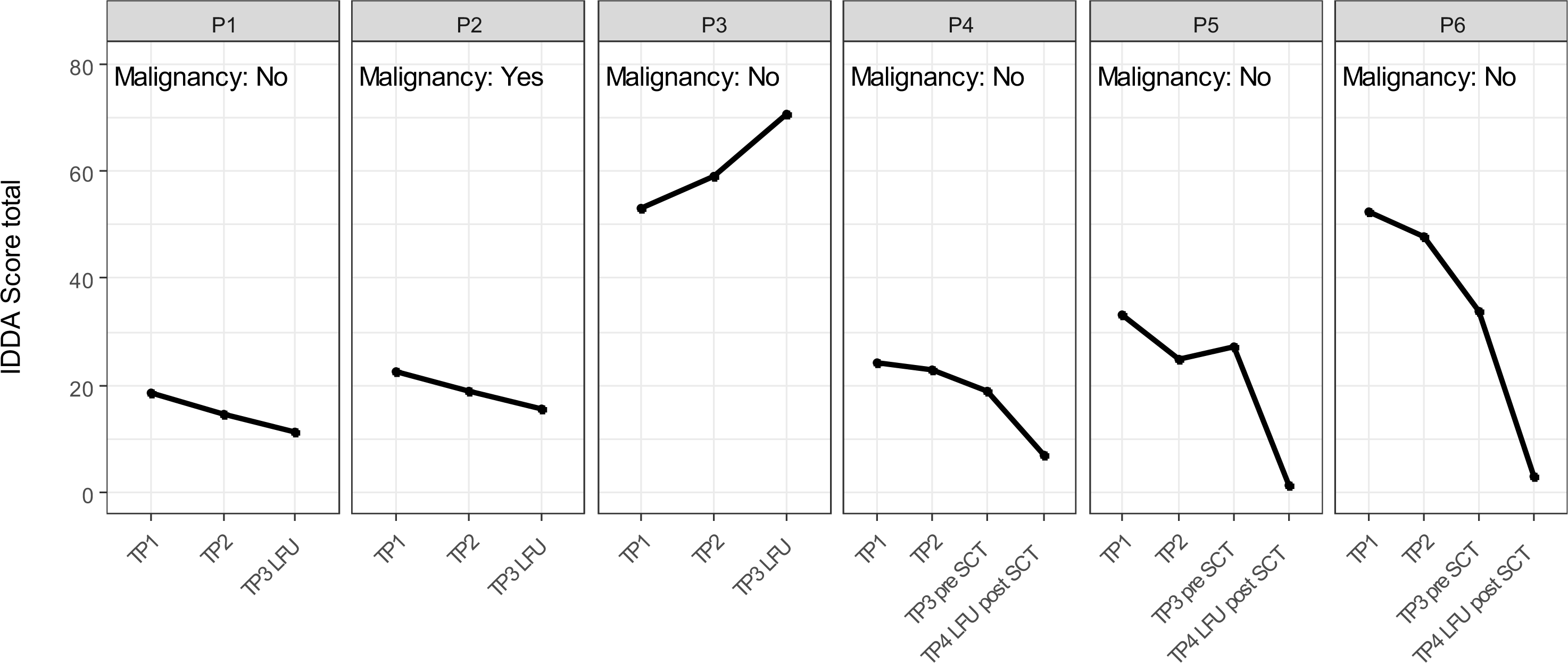
Development of the IDDA2.1 score under the therapy with abatacept and in P4-P6 post alloHSCT. The lower the score, the lower the severity of symptoms. TP1 marks the first measurement before starting abatacept therapy. TP2 and TP3 during abatacept and marks the last follow up for P1-P3 whereas P4-P6 have another TP4 after being transplanted. In P1 and P2 we can see a continuous slight improvement in disease activity. In contrast, P3 continued worsening the symptoms, despite partial response to abatacept. In P4-6, we see a significant reduction in disease activity after transplantation with scores in P5 and P6 almost being disease-free. IDDA, Immune deficiency and dysregulation activity score; TP, Time Point.

For three Patients alloHSCT was considered as the treatment of choice to achieve long-term stabilization of their immune phenotype. These patients showed a complete remission of autoimmune cytopenia and two of them no longer have any intestinal symptoms, these two also no longer show any failure to thrive. The third patient still has chronic diarrhea and weight stagnation. All three patients show a normalization of spleen size and a complete regression of the skin manifestations. Remaining in one patient is the pre-existing moderately controlled IDDM and another patient still has autoimmune thyroiditis at a stable level. The third transplanted patient had no previous endocrinopathies. None of the transplanted patients required further immunoglobulin replacement and there were no more chronic or recurrent infections, as two of the patients had before the transplant. (Figure 6B) While receiving abatacept four patients’ sIL2R levels decreased over time, correlating with the amelioration of their disease symptoms and/or treatment by alloHSCT. Two patients showed increasing levels (P3 and P4). P4 had residual disease symptoms post alloHSCT with a mixed chimerism currently under investigation.

**Figure 6B:**
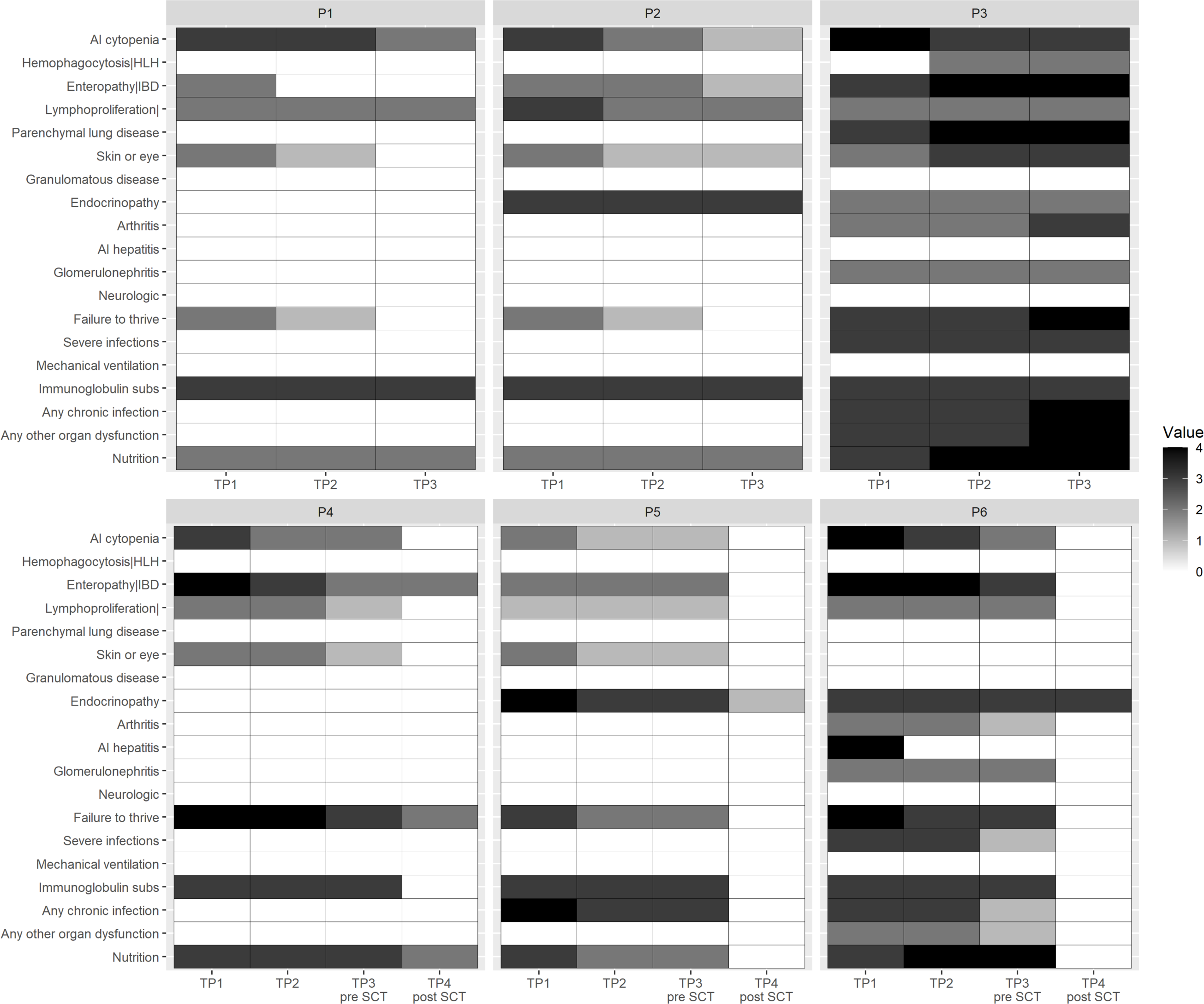
The IDDA2.1 score with each symptom at the respective measuring point. All 19 items (without malignancy, Karnovsky/Lansky Performance Scale and Hospitalization days) are shown for each patient. TP1 is the first measuring point without any therapy, TP2 is under abatacept therapy and TP3 before alloHSCT or at last follow up for non-transplanted patients and TP4 after alloHSCT. During biweekly intravenous administration of abatacept we observe a regression of symptoms, which is given by grey shading. P3 showed worsening of symptoms of enteropathy,GLILD and infectious complications. IDDA, Immune deficiency and dysregulation activity score; TP, Time Point.

## 4. Discussion

Maintaining a distinction between immunity and autoimmunity requires a complex interaction of activating and inhibiting factors. CTLA-4, mainly localized in intracellular vesicles of Treg, is a key regulator molecule in this cascade. By acting as an early negative checkpoint, CTLA-4 exerts a major influence on maintaining self-tolerance [28]. There is evidence that LRBA controls CTLA-4 expression by preventing its recycling in lysosomes, thereby supporting its trans-endocytic function and, as a result, the suppressive capacity of Treg. [19].

Treg compartment is a complex interactive mixture of different cell subsets. Gaining insight in Treg homeostasis by using further defining markers in addition to FoxP3 provides a ground for understanding complex diseases such as LRBA deficiency.

In this study we analyzed a cohort of six LRBA deficient patients with focus on their Treg subpopulations circulating in peripheral blood. Our data support the understanding that there is an age-dependent maturation of Treg subsets in healthy individuals, which is clearly visible in an increase of Helios^+^ natural Treg and memory Treg, while predominantly naïve Treg decrease by the age of 40 years. With focus on LRBA deficient individuals, we observe that this maturation is severely affected resulting in very low levels of circulating Helios^+^ natural Treg, CD45RA^+^ naïve Treg, FoxP3^hi^ effector Treg and CD25^hi^CD127^low^ Treg. Interestingly, there is a shift towards more memory Treg in these patients. All the patients of our cohort showed normal values for Helios^-^ induced Treg. The positive therapeutic effect of abatacept has already been shown in different case series. [29] [30] Our goal was to follow the Treg subsets during the treatment. We observed a unique pattern in Helios^+^ natural Treg as there was a decrease in numbers shortly after abatacept treatment start, followed by a significant increase, up to normal range, over the course of the disease. Clinical improvement in our patients confirmed the beneficial effect of abatacept, which was also accompanied by decreasing levels of sIL2R. Nevertheless, for three patients the alloHSCT was performed to achieve long-term remission.

Our data present a detailed phenotyping of the circulating Treg in peripheral blood. Given the fact that there is growing evidence on existence and function of tissue resident Treg [31], further studies in LRBA deficient individuals or disease models are needed to explore the role of tissue Treg subsets in affected organs.

Clinical scoring systems aim to objectify the disease severity. While for a mainly pediatric cohort, the CHAI-score was not feasible, the IDDA2.1 score was found to be a feasible and reliable scoring system to assess the disease severity in the beginning and over the course of treatment.

## 5. Conclusion

In summary, we provide information on the Treg profile of peripheral blood samples from a mainly pediatric cohort with LRBA deficiency compared to that of healthy age-matched cohorts. Helios^+^ natural Treg, naïve, effector and CD25^hi^CD127^low^ Treg were severely lacking. Only natural Treg increased with a unique pattern to normal levels during abatacept treatment. Clinical scoring systems such as IDDA 2.1 score can be helpful to assess and unify the evaluation of disease severity.

## Acknowledgments

We thank all patients, family members and physicians who participated in this study and the healthy individuals for blood donation. We especially thank technicians of the Laboratory for Stem Cell Transplantation and Immunotherapy of the Children’s Hospital Frankfurt.

## 7. Author contributions

SB, SD, ESM conceptualization, data analysis, investigation, writing. SH supervised flowcytometry assays. CK provided samples of healthy controls. PB, LM provided data and revised the manuscript. SD, LML, JF performed experiments and analyzed the data.

---

**Suppl. Figure 1:**
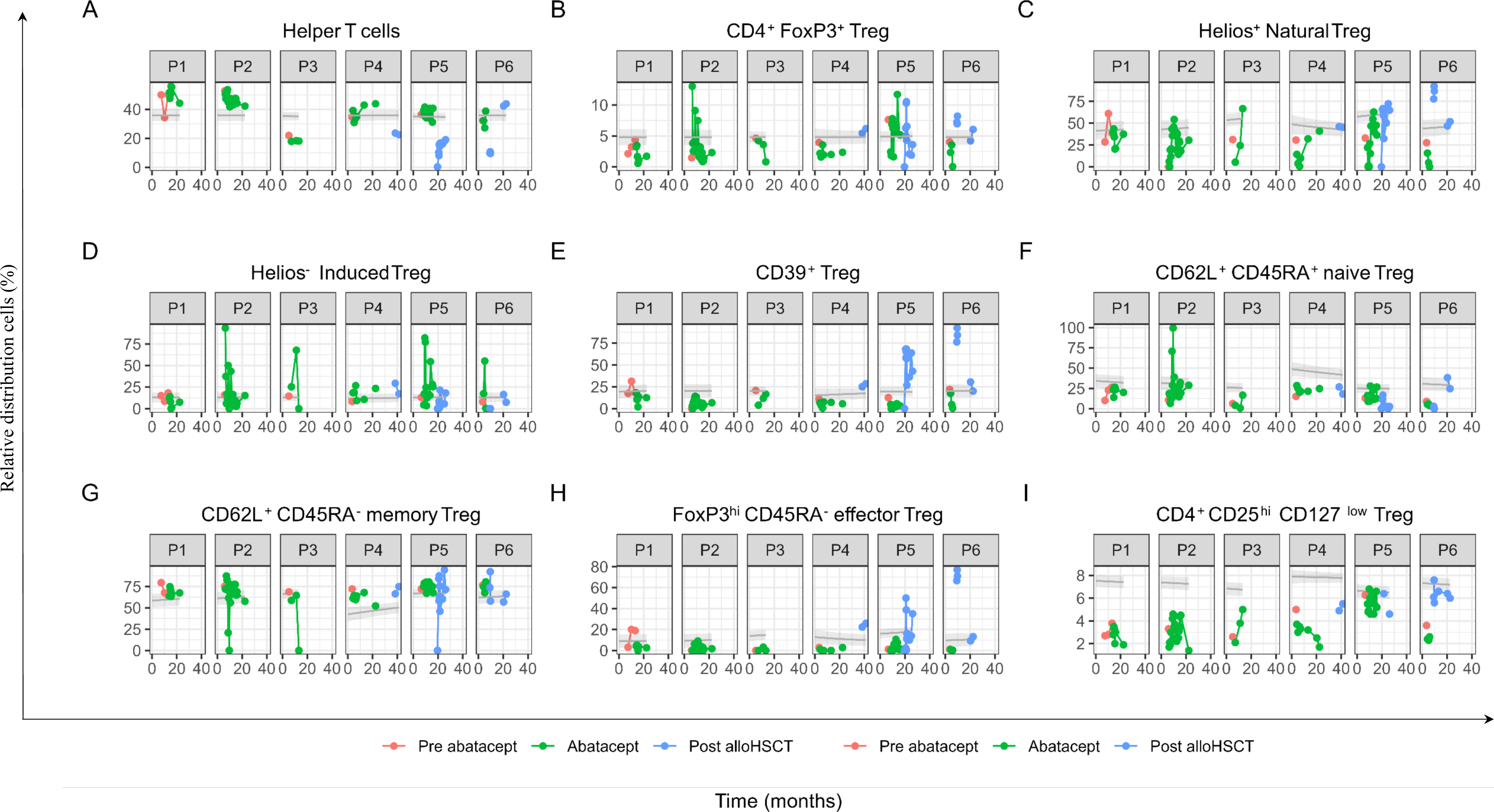
Longitudinal Data analysis from six patients suffering from LRBA Deficiency during intravenous abatacept administration and after performance of alloHSCT in P4-P6. Each dot represents a measurement time point, with red dots marking measurements before starting abatacept therapy, green dots during biweekly intravenous abatacept therapy and blue dots in P4-P6 after alloHSCT. Abatacept therapy was stopped after receiving alloHSCT. On the y-axis, the values are shown as the percentage of lymphocytes (for A) and T helper cells (for B) or CD25^hi^ Treg (for C-I). The x-axis shows the period of observation in months, starting with the time of the first measurement. Reference values from healthy volunteers are represented with the grey shaded area. alloHSCT, Allogeneic Haematopoietic Stem Cell Transplantation.

**Suppl. Table 1:**
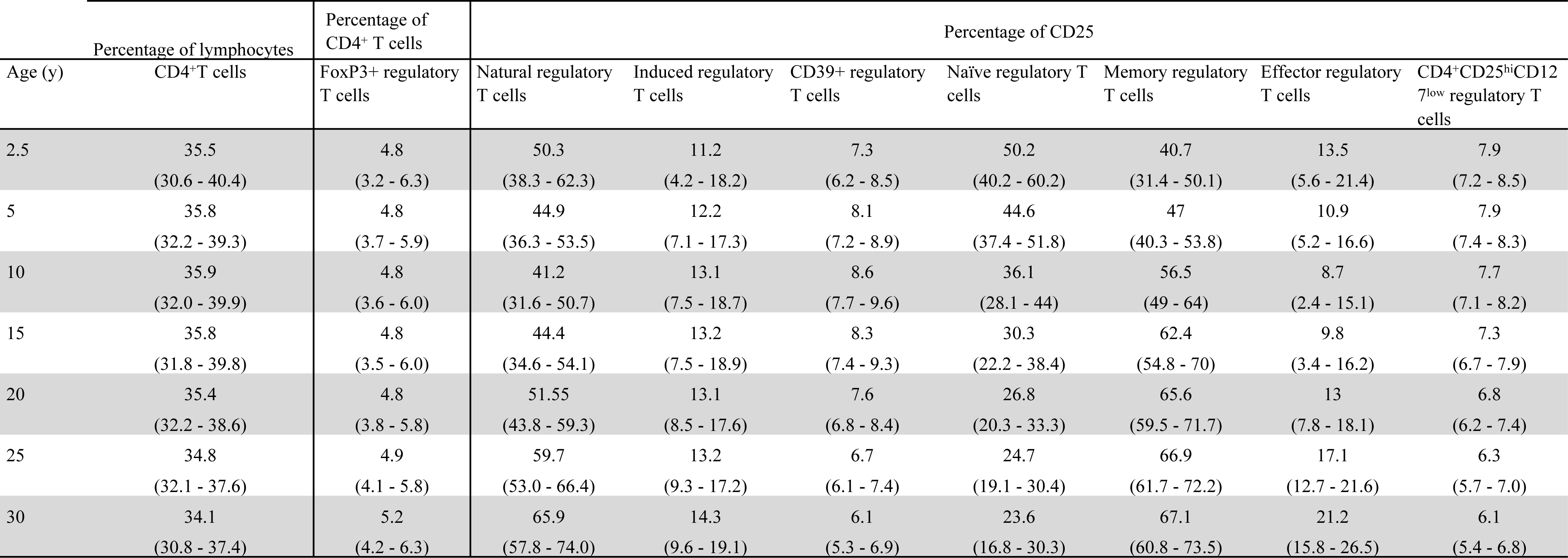
The table presents the predicted reference of relative distribution cells (%) from our B-spline linear model with 95% confidence interval according to age in parenthesis. Relative frequencies (% of CD4^+^ T helper) for FoxP3^+^ regulatoty T cells and for each Treg subpopulation (% of CD4^+^CD25^hi^) of healthy donors given for ages of 2.5, 5, 10, 15, 20, 25 and 30 years. The total amount of FoxP3 Treg remains at about 5%, while there is a gradual increase in natural Treg, induced Treg, memory and effector Memory Treg by the age of 40. Naïve Treg decrease over time. FoxP3, Forkhead-Box-Protein P3.

## Abbreviations

AI: Autoimmunity
AlloHSCT: Allogeneic haematopoietic stem cell transplantation
APC: Antigen presenting cell
CD39: Ectonucleoside triphosphate diphosphohydrolase-1
CHAI: CTLA-4 haploinsufficiency with autoimmune infiltration (CHAI)
CTLA-4: Cytotoxic T-lymphocyte associated protein-4 (CD152)
FCS: Fetal Calf Serum
FoxP3: Forkhead box protein P3
HLH: Hemophagocytic lymphohistiocytosis
IDDA: Immune Deficiency and Dysregulation Activity
IDDM: Insulin dependent diabetes mellitus
IL: Interleukin
IPEX: Immunodysregulation polyendocrinopathy enteropathy X-linked
I.v.: Intravenous
LAG-3: Lymphocyte activation gene-3
LRBA: Lipopolysaccharide-responsive beige-like anchor protein
OX40: Tumor necrosis factor receptor superfamily, member 4 (CD134)
PBS: Phosphate buffered saline
SD: Standard deviation
sIL2R: Soluble interleukin-2 receptor
Treg: Regulatory T cell
4-1BB: Tumor necrosis factor ligand superfamily member 9 (CD137L)

